# High resolution *Streptococcus pyogenes* core genome MLST and LIN coding scheme for outbreak detection

**DOI:** 10.64898/2026.07.10.737715

**Authors:** Yan Ryan, Keith A Jolley, Henry Hearn, Kasia M Parfitt, Steven Platt, Theresa Lamagni, Kartyk Moganeradj

## Abstract

*Streptococcus pyogenes* is a globally important pathogen responsible for at least 500,000 deaths a year, causing significant burden on healthcare systems. It is the causative agent for ailments such as impetigo and strep throat to septicaemia and necrotizing fasciitis. Assessment of genetic relatedness for the detection of outbreaks within communities or healthcare facilities is vital in decreasing the propagation of *S. pyogenes* within these settings, alongside epidemiological data. As the volume of isolates being sequenced increases year on year, more scalable and sharable methodologies of assessing genetic relatedness are required by reference laboratories and for international collaboration. LIN codes, applied to core genome MLST (cgMLST) represent a method which is extensible to large scale whole genome sequencing (WGS) while still being sufficiently sensitive to detect outbreak clusters. Here we present a novel cgMLST and LIN code scheme, hosted by PubMLST, enabling international collaboration and global tracking of variants, that is highly scalable and usable for all. The schemes are available at https://pubmlst.org/organisms/streptococcus-pyogenes.

**Data Summary:** Genome sequences and metadata are available at https://pubmlst.org/organisms/streptococcus-pyogenes. PubMLST and ENA accessions and metadata can additionally be found in the supplementary data. Raw reads for UKHSA sequences are available in ENA study PRJEB115996.

**Impact Statement:** Streptococcus pyogenes is a globally relevant pathogen capable of causing invasive and non-invasive disease across a multitude of settings. Assessment of genetic relatedness is an increasingly important aspect of managing outbreaks, requiring solutions that are scalable, high resolution and comparable across laboratories. Here we present a high resolution core genome multi locus sequence typing (MLST) and associated life identification number (LIN) code scheme, The schemes were developed using a combination of 4,916 UKHSA and 2,391 publicly available S. pyogenes isolates in order to cover a wide range of EMM types both within the UK and globally. These new schemes enable high resolution typing of S. pyogenes isolates, suitable for analysis of lineages to genomic epidemiology in outbreak detection and management. Both cgMLST and LIN code schemes are available on PubMLST as an open access resource for the public health and academic communities and can enable both intra laboratory and global coordination.

## Introduction

Group A Streptococcus (GAS), known formally as *Streptococcus pyogenes*, remains a source of significant mortality, with an estimated 306,000 deaths associated with rheumatic heart disease in 2019 (Roth, Mensah et al. 2020) and invasive GAS (iGAS) resulting in a mortality rate of up to 45% (Craik, Hla et al. 2022), causing considerable burden to healthcare systems across the resource spectrum (Thacharodi, Hassan et al. 2025). GAS can cause a range of ailments, ranging in severity from sore throat and impetigo to necrotizing fasciitis, streptococcal toxic shock-like syndrome and septicaemia (Brouwer, Rivera-Hernandez et al. 2023). Cluster-based transmission can be an important factor for GAS, in one study accounting for 64% of events (Metcalf, Nanduri et al. 2022), highlighting the importance of determining the genetic relatedness of isolate clusters. Since 2019 in the United Kingdom, the number of detected outbreaks and cases of iGAS have increased, with 13% of isolates identified as part of a cluster, the majority of which occurring in hospitals, care homes and homeless shelters (Smith, Marchant et al. 2025).

*S. pyogenes* creates an important cell surface protein known as the M protein (EMM type), which is a virulence factor involved in adhesion, colonisation and anti-phagocytosis (Oehmcke, Shannon et al. 2010). The EMM gene is based in the mgA regulon, and has considerable diversity, with over 200 EMM genes and their translated M proteins currently recognised (Bessen and Lizano 2010). EMM genes are categorised into the EMM clusters A-E, dependent on mgA regulon structure with EMM clusters displaying preferential tissue tropisms (Bessen and Lizano 2010). This ubiquity of M proteins has made it a promising vaccine candidate, with the M protein containing conserved regions for binding to host proteins, such as a C4b-binding protein region representing a possible vaccine target (Wang, Kuliyev et al. 2023). These conserved regions have shown promise as cross opsonising multi-valent vaccine targets, with multi-valent vaccines covering a broad variety of M types (Frost, Laho et al. 2017, Ghosh 2018). EMM type diversity is inversely correlated with Human Development Index (HDI), limited in high income nations, increasing in diversity as HDI decreases, with 15 EMM-clusters representing 95% of global strain diversity (Smeesters, deCrombrugghe et al. 2024). The M protein has historically been used for determination of species by Lancefield testing (Lancefield 1933), followed by serological Polymerase Chain Reaction (PCR) based (Beall, Facklam et al. 1996) and WGS based typing of the EMM gene (Kapatai, Coelho et al. 2017). EMM typing provides useful epidemiological information at both local and global levels for tracking outbreaks and trends over time.

While EMM typing provides a useful marker for GAS identification, it’s resolution for outbreaks is limited for effective delineation. No universally comparable method of genetic relatedness has been available, with different national reference laboratories utilising a range of solutions from Single Nucleotide Polymorphism (SNP) based (Davison, Clements et al. 2025) to core genome multi locus sequence typing (cgMLST) allelic distances (Beres, Olsen et al. 2024). SNP derived schemes are effective locally but challenging to share without the same reference sequence and SNP-calling pipeline, whereas cgMLST schemes enable easier sharing of sequence types (STs) (Maiden, vanRensburg et al. 2013). However, this information does not immediately elucidate genomic relatedness without determining the distances between the allelic profile for each ST. cgMLST indexes the allelic variation across core genes, with each unique combination of alleles assigned a cgST number. cgMLST can deflate the distance between alleles compared to SNP based analysis, as one or more SNP differences is only counted as a distance of one, whereas a SNP based analysis would reflect the total sum of SNP differences between alleles. However, SNP-based analysis can be skewed by the presence of mobile genetic elements, where the existence of one can inflate the distance between two isolates by the contents of the added gene. Whereas a cgMLST based scheme would only record a difference of one.

LIN® (Trademark registered by This Genomic Life, Inc, Floyd, VA, USA) (Life Identification Number) codes, initially envisioned using average nucleotide identity, provide a stable nomenclature to cluster profiles at differing thresholds of identity (Vinatzer, Tian et al. 2017). Later implementations enabled the use of species specific cgMLST schemes for LIN code assignment (Hennart, Guglielmini et al. 2022). LIN codes take a cgMLST profile, assign it to a series of thresholds, dictated by the number of shared genes, ranging from broad species structure, to an identical cgMLST profile that may differ only at loci missing in one of the profiles; each new profile within a threshold produces an increment in that level, creating a stable, and expansible “LIN code” that is readily comparable (Hennart, Guglielmini et al. 2022). These LIN codes can be readily stored in a database such as PubMLST (pubmlst.org). LIN codes are read from left to right, increasing in relatedness with each threshold.

This paper showcases the development and use of a cgMLST derived LIN code scheme for *S. pyogenes* that has been made publicly available and its utility for assessing genetic relatedness for global population dynamics and public health outbreak investigations.

## Methods

### Isolate selection

In order to cover as many of EMM types as possible, UKHSA isolates (n=4916) were supplemented with an additional tranche of isolates downloaded from NCBI (n=2391) (Supplementary table 1). UKHSA isolates spanned from 2023 to 2025 for typing and assessment of genetic relatedness for outbreak control. The Bacteriology reference department (BRD) user manual (https://www.gov.uk/government/publications/bacteriology-reference-department-brd-user-manual) requests all iGAS isolates identified in England to be sent to UKHSA, thus is a representative collection of sporadic and outbreak strains.

NCBI isolates were selected from publicly available, annotated *S. pyogenes* genomes with no selection based on sample date or geography and downloaded in May 2025.

EMM_typer (https://github.com/MDU-PHL/EMMtyper) was used to determine the presence of 152 different EMM types excluding subtypes among this dataset, using the latest EMM database from the CDC (https://www2.cdc.gov/vaccines/biotech/strepblast.asp).

### Sequencing

Sequencing of UKHSA isolates (n=4916) was performed using an Illumina Nextera XT library prep kit on an Illumina Nextseq 1000 with a read length of 2×100 base pairs. FASTQ files were trimmed and selected for quality scores of Q30 or better by Trimmomatic (version 0.32) (Bolger, Lohse et al. 2014).

### Phylogenetic analysis

SNP based phylogenetic analysis was performed as described by Chalker et al (Chalker, Jironkin et al. 2017). PHEnix (https://github.com/ukhsa-collaboration/PHEnix) (version 1.4.3) utilised BWA-mem (Li 2013) (version 0.7.12-r1039) to map reads against a reference of the appropriate EMM type for each EMM type (Supplementary Table 2), with GATK (version 2.6.5) used to call SNPs. Gubbins (Croucher, Page et al. 2015) (version 3.4.3) was used to remove areas of recombination and create a maximum likelihood tree using (RAxML-NG v. 1.2.2)(Croucher, Page et al. 2015). The resulting tree was visualised in R using ggTree (version 3.14.0) (Xu, Li et al. 2022)

A cgMLST profile-based minimum-spanning analysis was performed using GrapeTree (Zhou, Alikhan et al. 2018) as hosted on PubMLST (Jolley, Bray et al. 2018) using the MSTreeV2 option for tree generation.

### cgMLST scheme and allele calling

ChewBBACA (Silva, Machado et al. 2018); (version 3.3.10), was used to call alleles of the above datasets for both the ChewieNS and Toroop schemes.

ChewBBACA’s ExtractCgMLST option was then used to create a cgMLST scheme, with loci present in 98% of isolates from the above isolate dataset and ChewieNS wgMLST (whole genome MLST) scheme and Toroop cgMLST scheme.

The ChewieNS scheme was further curated to remove inconsistent, non-conserved start sites and alternate length alleles, and was evaluated against the un-curated and Toroop scheme by assessing correlation with a matched SNP distance matrix using the Mantel test, with the highest scoring scheme (Manually curated ChewieNS) being taken forward.

### Mantel test

Mantel tests were performed using R (version 4.4.0) and the function mantel() from the package vegan (2.6-4)(Jari Oksanen; Gavin L. Simpson; F. Guillaume Blanchet; Roel 2025) to compare matrices generated from SNP distances and cgMLST allelic distances.

SNP matrices were created as previously described (Dolan, Coelho et al. 2025). Variant data post Gubbins removing areas of recombination were used as input to create a distance matrix from snp-dists (version 0.8.2 https://github.com/tseemann/snp-dists).

The cgmlst-dists (https://github.com/tseemann/cgmlst-dists) (version 0.4.0) software was used to determine the distance of the allelic profiles as determined by the ChewBBACA allele_call module.

### LIN codes

Adjusted rand indices were calculated using the R package fossil (version 0.4) (Vavrek 2011)

MSTClust (https://gitlab.pasteur.fr/GIPhy/MSTclust, version 0.21b) was used to generate silhouette scores to identify cluster cohesiveness.

LIN codes were derived from the final cgMLST scheme using nine thresholds: 528;285;124;25;10;5;2;1;0 allelic differences, using the LIN code module on BIGSdb. The scheme is hosted on https://pubmlst.org/organisms/streptococcus-pyogenes.

The resulting LIN codes were assessed against a range of EMM types, both clonal and non-clonal to cover a representative sample of the *S. pyogenes* population. Clusters were assigned to isolates within previous outbreaks based on SNP distances and LIN codes.

Isolates with 10 or less SNPs were included in the same cluster, and isolates matching at the 5-10 allele threshold were linked to the same cluster.

## Results

### cgMLST scheme assessment

The initial cgMLST schemes were defined by using both NCBI and UKHSA *S. pyogenes* isolates (n=7307) to determine a 98% presence cgMLST scheme, firstly by allele calling with ChewBBACA against the S. pyogenes Chewie-NS wgMLST scheme (i), and the Toroop *S. pyogenes* scheme (ii). These allelic profiles were then used to generate cgMLST schemes with allele presence in a minimum of 98% of isolates using the ChewBBACA extract cgMLST tool. These new 98% cgMLST schemes were then taken forward for accuracy assessment.

To assess the accuracy of the cgMLST schemes, isolates representing 10 EMM genes were selected (Table 1). This included both clonal and non-clonal EMM types, with a minimum of 25 isolates per EMM type. The Mantel test was used to calculate a score of the statistical correlation between SNP distances and allelic distances for the same isolates. Some discrepancy was expected due to the differing methodologies as the SNP distances have masked areas of recombination in the genome. Whereas MLST approaches include these regions but help mitigate the effects of recombination by counting any allelic difference the same, irrespective of the number of nucleotide differences.

**Table 1.** Table of Mantel test scores, comparing cgMLST matrices against SNP matrices by EMM type, showing clonality and count of isolates per EMM type.

| EMM type | Isolate<br>count<br>(n) | Clonality | Chewie NS<br>(i) | Toroop<br>(ii) | Chewie<br>NS<br>Curated<br>(iii) |
| --- | --- | --- | --- | --- | --- |
| 1 | 243 | Clonal | 0.9063 | 0.8644 | 0.8936 |
| 3 | 187 | Clonal | 0.7967 | 0.7162 | 0.8322 |
| 6 | 248 | Clonal | 0.9676 | 0.9314 | 0.9714 |
| 8 | 66 | Clonal | 0.8573 | 0.8325 | 0.8873 |
| 59 | 72 | Clonal | 0.7632 | 0.7439 | 0.7659 |
| 33 | 52 | Non-Clonal | 0.9219 | 0.8946 | 0.9251 |
| 28 | 60 | Non-Clonal | 0.9313 | 0.93 | 0.9331 |
| 77 | 172 | Non-Clonal | 0.8656 | 0.8647 | 0.8972 |
| 82 | 26 | Non-Clonal | 0.8479 | 0.8552 | 0.8525 |
| 11 | 189 | Non-Clonal | 0.9646 | 0.9494 | 0.9717 |
| Average Mantel Score |  |  | 0.8824 | 0.8582 | 0.893 |

**Table 2.** Table summarising cutoffs used for S. pyogenes LIN coding scheme by percentage of cgMLST scheme, number of allelic differences, and the Mean of nAUC Silhouette and nAUC Wallace scores.

| LINcode threshold | Cutoff % Allelic differences | Cutoff allelic differences (n) | Mean of nAUC silhouette and nAUC Wallace |
| --- | --- | --- | --- |
| 1 | 55.93 | 528 | 0.96 |
| 2 | 76.21 | 285 | 0.97 |
| 3 | 89.65 | 124 | 0.95 |
| 4 | 97.91 | 25 | 0.85 |
| 5 | 99.17 | 10 | 0.85 |
| 6 | 99.58 | 5 | 0.91 |
| 7 | 99.84 | 2 | 0.85 |
| 8 | 99.92 | 1 | 0.85 |
| 9 | 100 | 0 |  |

The Mantel test showed variance across all EMM types and cgMLST schemes, with the Chewie-NS (Friães, Mamede et al. 2022) (https://chewbbaca.online/species/1) scheme providing the highest average score (Table 1). Further investigation of the Chewie-NS scheme revealed loci with a number of issues, such as inconsistent start sites, start codons, and differing gene lengths. A curated ChewieNS scheme (iii) was created which resolved the previous issues. After re-running the analysis, Mantel test results for the curated ChewieNS scheme, showed an improvement over the original scheme of 0.0106. As the curated Chewie-NS scheme had the best score it was selected for use as the S. pyogenes cgMLST scheme to be hosted on PubMLST and for developing the LIN coding scheme.

EMM 59 shows the lowest correlation between the SNP matrix and cgMLST matrices, with an average score of 0.758. EMM 11 has the highest correlation with an average score of 0.9619. EMM59 exhibits a lower correlation due to nonlinear scaling between the number of SNP differences and allelic differences, with ranges of 10 SNPs resulting in 2-3 allelic differences. EMM59 may have a small number of genes with a high selective pressure, increasing mutation among these genes, or the regions under selective pressure are not part of the cgMLST scheme but present for SNP calling. Non-clonal isolates show a greater correlation between cgMLST and SNP (clonal mean 0.87008 compared to a non-clonal mean of 0.91592).

### LIN code threshold determination

LIN code thresholds are determined by a percentage of shared alleles and can range from 0-100% allelic mismatches. Assessment of pairwise allelic dissimilarities can produce natural peaks and troughs representing structures within a bacterial population. These can be used to capture the structure within LIN code thresholds. Visualisation of this with pairwise allelic distances showed primarily a single peak at 1075 allelic differences. There were also several smaller peaks observed between 0 and 50 allelic differences (Figure 1). No pairwise mismatches were greater than 1139 out of 1198 genes within the cgMLST scheme. This indicated that isolates shared a minimum of 59 loci or 4.9% with at least one other isolate in the dataset.

**Figure 1.**
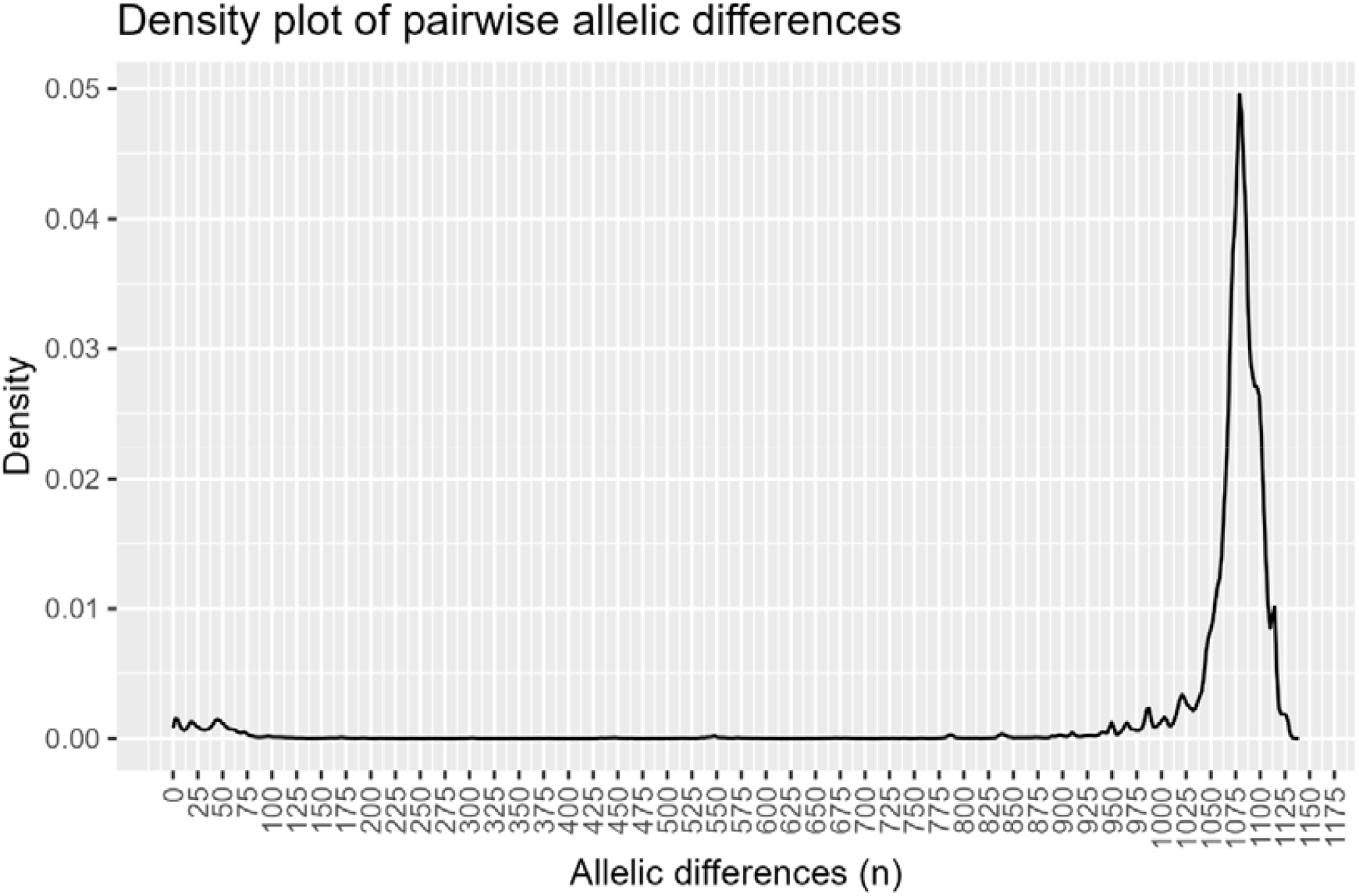
Density plot of S. pyogenes pairwise allelic differences, showing a primary peak around 1075 allelic differences, with some minor peaks between 0-75 allelic differences.

Relatively little structure can be observed from the pairwise allelic differences, other than the primary peak at 1075 allelic differences. Therefore, ridgeline plots (Wilke 2025) composed of EMM types (Figure 2), and STs (Figure 3) can be used to determine appropriate LIN code thresholds through the delineation of samples. Concatenation of the STs with EMM type compared to ST alone showed a decrease in the maximum number of allelic differences from ∼900 to 500. Both plots showed most density peaks between 0-100 allelic differences, with a small subset of differences at 100-200 allelic differences, and EMM 11 as an outlier with up to 900 allelic differences (Figure 3). These coincide with the minor peaks seen in Figure 1 in the all-by-all allelic difference density plot, at the 0-100 allelic difference range. These plots indicate that some potentially viable cut offs should be around the 100 allelic difference, 200 allelic differences and over 900 allelic difference range. The addition of EMM types to ST removed more distantly related isolates with shared MLST profiles, as evidenced by EMM:ST having a lower maximum number of identified allelic differences (Figures 2 and 3).

**Figure 2.**
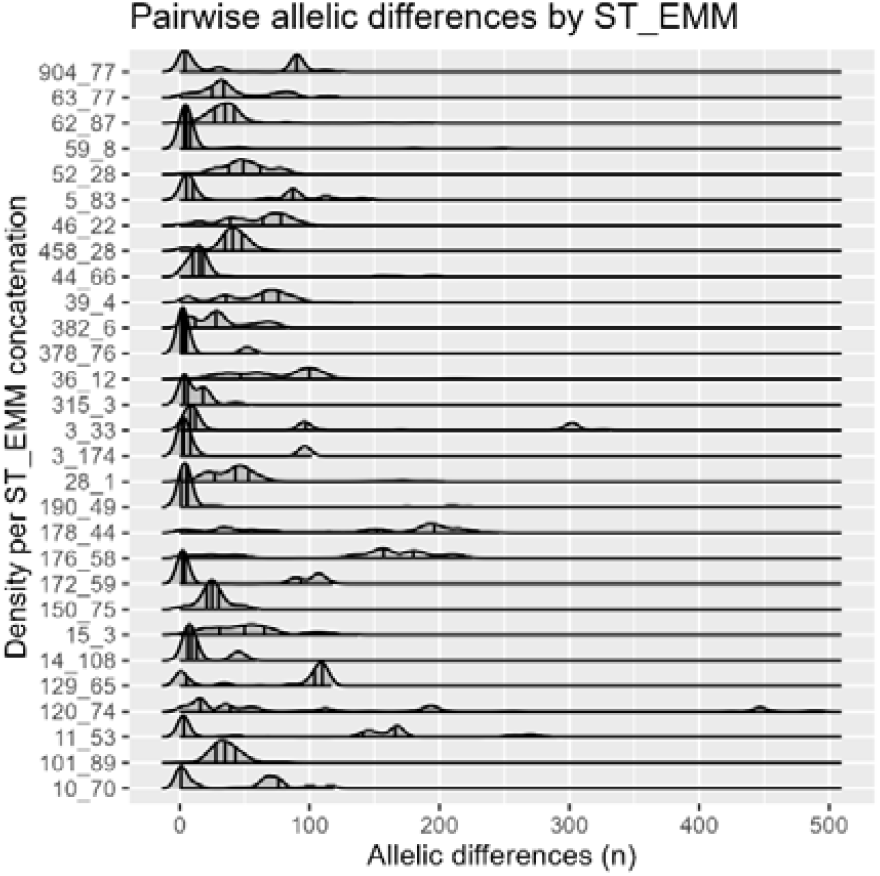
Ridge plot of sequence type (ST) and EMM type displaying density of allelic differences. The majority of pairwise allelic differences occur under 100 allelic differences, with a smaller subset having peaks in the 100-200 differences range, with ST3_EMM33 having one of several peaks at 300 differences, and ST12_EMM74 likewise having a peak at ∼450 allelic differences.

**Figure 3.**
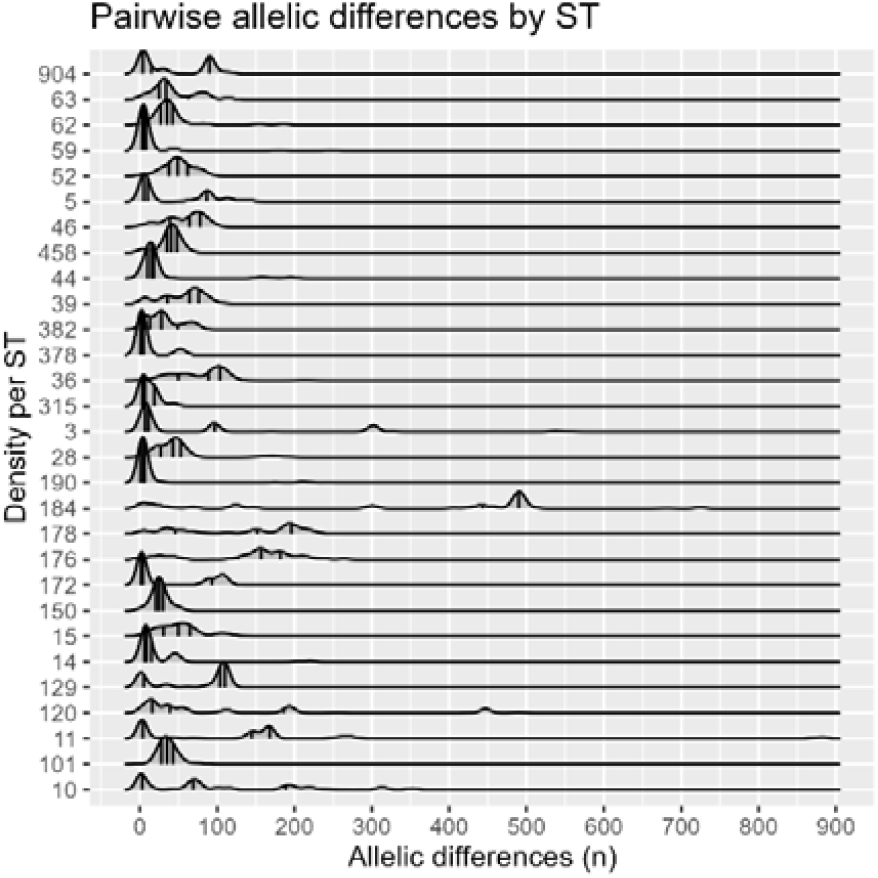
Ridge plot of S. pyogenes sequence type displaying density of allelic differences. The majority of allelic differences peak at under 100 differences, with ST184 being a clear outlier at 500, and ST11 having a minor peak around 850 differences. Several show multiple peaks indicating a high level of intra ST diversity.

As the all versus all density plot and ridge plots are respectivelly head and tail heavy, leaving little additional information for cutoff thresholds, MSTclust (https://gitlab.pasteur.fr/GIPhy/MSTclust) was used to ascertain additional optimum cutoff levels for LIN code thresholds. MSTclust was used to create a minimum spanning tree from the previously created cgMLST profile, and then analysed to identify optimum points by MST estimated statistics.

For this analysis the mean nAUC (naÏve area under curve) Wallace and nAUC silhouette scores were plotted (Figure 4) to further identify optimum cutoff values. The silhouette index indicates clustering consistency within a dataset and Wallace coefficient indicates agreements within groups of clusters. Therefore a higher mean score of both the Wallace coefficient and Silhouette Index could be plotted to identify optimum cutoffs. At 1 allelic difference (0.08%) the mean nAUC Wallace/Silhouette score was 0.77/1, rising rapidly to a peak of 0.97/1 at 285 (23.7%) allelic differences, before a rapid decline from 910 allelic differences (76%). Therefore the most optimum cutoffs lie between 5% and 76% allelic differences.

**Figure 4.**
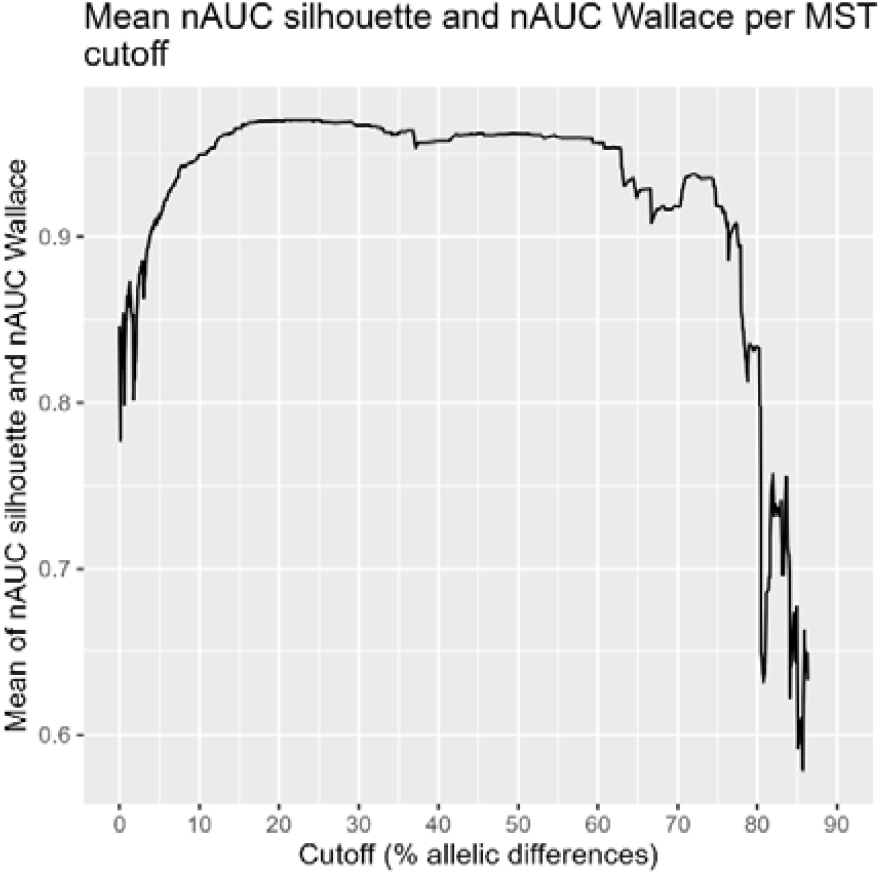
Line plot of mean of nAUC silhouette and nAUC Wallace by gene cut off, as calculated by MST. Higher values indicate better clustering consistency and agreement within clusters.

**Figure 5.**
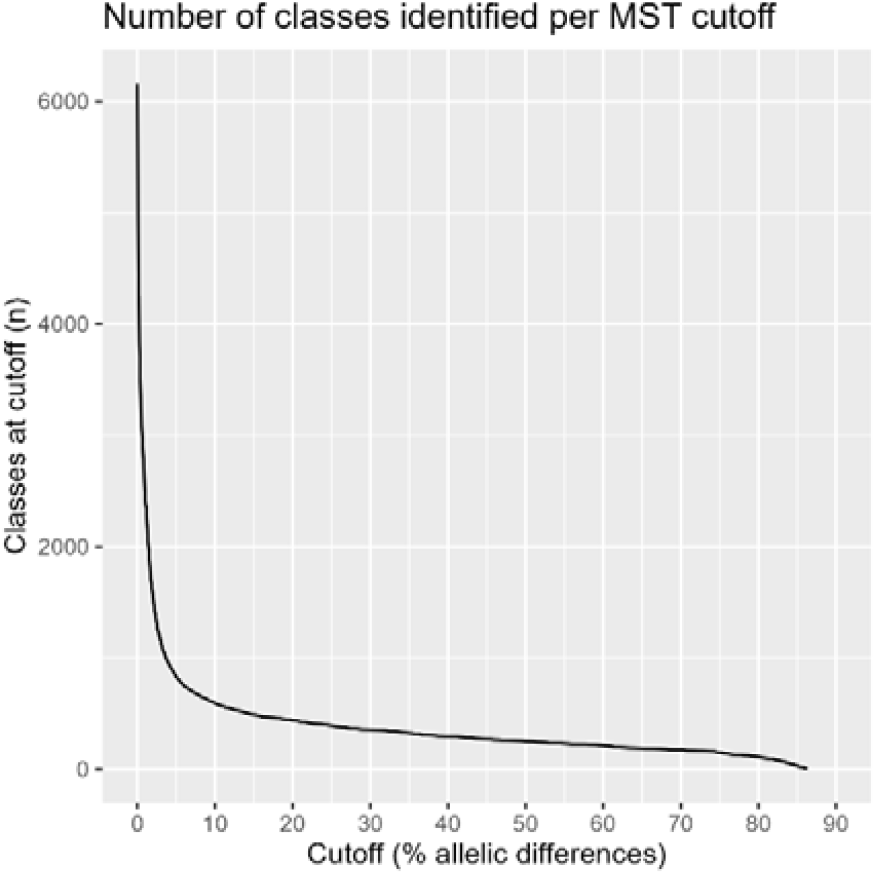
Number of classes identified by MST at each gene cutoff. Shows a rapid decline in the number of clusters as the gene cutoff increases, reaching 0 classes at 96%.

Therefore, the following 9 thresholds were chosen as the allelic difference thresholds for the *S. pyogenes* LIN code scheme; 528, 285, 124, 25, 10, 5, 2, 1 and 0. The cutoffs at 528, 285 and 124 alleles represent differences in population structure, with 25, 10, 5, 2, 1 and 0 providing high granularity discrimination for outbreak investigations and are concordant with other LIN code schemes.

The ability of LIN codes to match to both EMM type and ST was undertaken both statistically (Table 3) and visually by annotating the output of the GrapeTree (Zhou, Alikhan et al. 2018) plugin on PubMLST. The minimum spanning tree was annotated with EMM, MLST ST and LIN code threshold 1 (Figure 6). Separation of multi-lineage EMM types can be seen in Figure 6, with strains such as EMM89 (dark brown at 2 o’clock and 9 o’clock on the clock face (Figure 6)) showing clear distinction as having branched at the EMM3 root, being identified as ST 101 and LIN 0: 215 for the most common EMM89 lineage at 2’oclock, and the smaller population at 9 o’clock, with ST 380 and LIN code threshold 248 (data not shown on figure, see Supplementary Table 1). Other EMM types such as EMM49, EMM4 and EMM77, show a predominant strain, correlating with broadly with a single ST type and LINcode 0 threshold, along with several smaller populations, with a lack of visualised ST or LIN code threshold 0 designation due to their rarity.

**Table 3.** Listing breakdown of first four LIN coding thresholds, the number of loci used per cutoff, and the adjusted Rand scores when matched against EMM type and MLST ST.

| LIN code thresholds | Adjusted Rand EMM type | Adjusted Rand MLST ST |
| --- | --- | --- |
| 1 | 0.881 | 0.849 |
| 1_2 | 0.941 | 0.918 |
| 1_2_3 | 0.932 | 0.912 |
| 1_2_3_4 | 0.849 | 0.869 |
| 1_2_3_4_5 | 0.737 | 0.786 |

**Figure 6.**
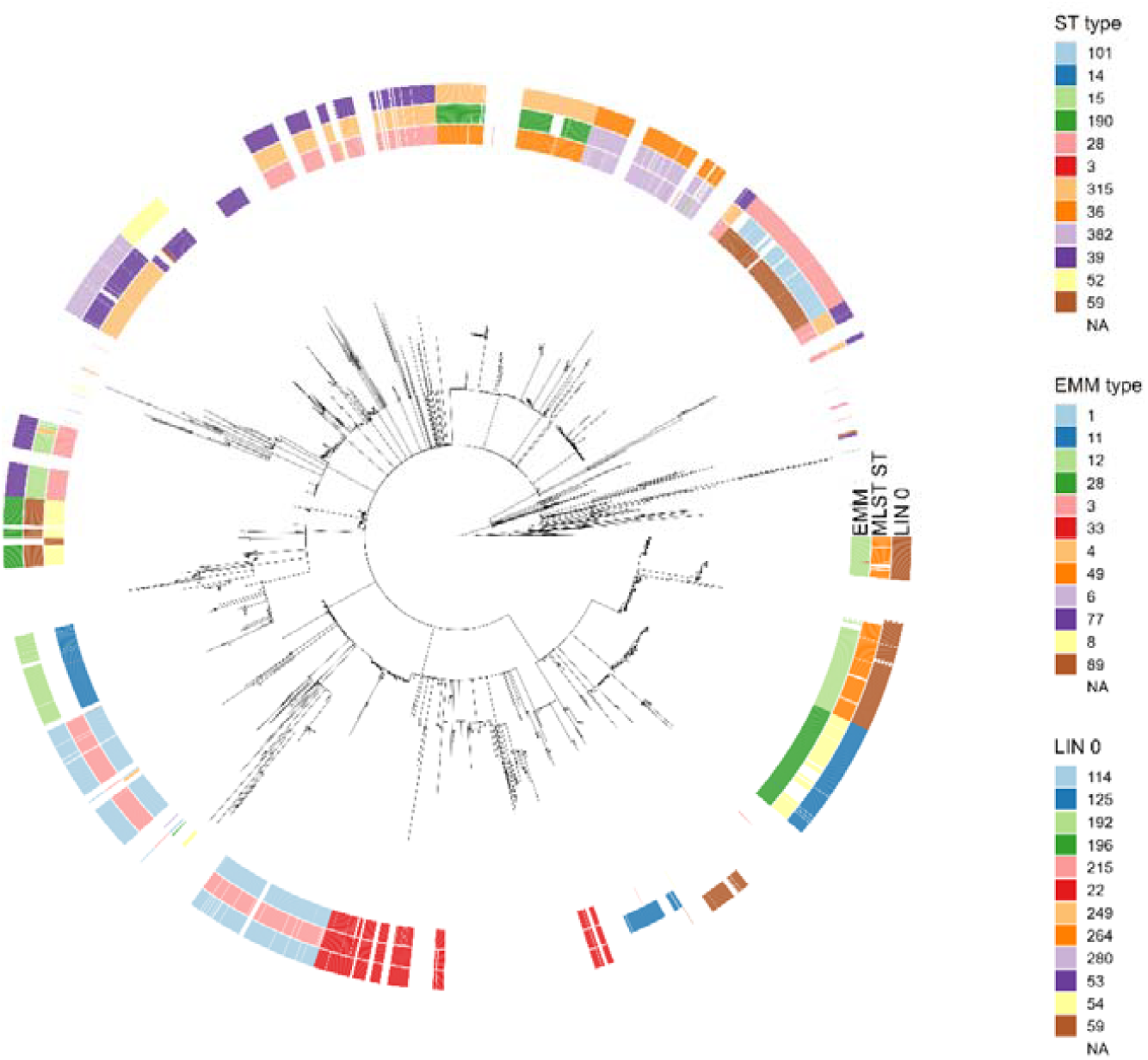
Phylogenetic tree of all test isolates submitted to PubMLST for development and testing of this scheme, showing cgMLST derived phylogenetic structure as created by GrapeTree using the MSTreeV2 option and visualised in ggTree. Top 12 categories for EMM type, ST and LIN threshold 0 are highlighted, with all others left blank. Visually confirms the high concordance as shown in Table 3. between EMM type, ST and LIN threshold 0.

The Rand index is used to assess the correlation between similarity of data clusters, with the adjusted Rand index (aARI) correcting for chance, where 0 indicates no correlation and 1 perfect correlation. The aARI scores for LIN codes to both EMM type and MLST ST show a high measure of similarity, peaking 0.941 and 0.918 respectively at the second LIN code level, decreasing to 0.932 and 0.912 respectively third level, and decreasing as the additional levels add granularity and therefore an increase in clusters.

Analysis of LIN code suitability as compared to known multi lineage strains and current EMM and ST typing shows good comparability and effective delineation of S. pyogenes species specific genomic structure, while avoiding the potential pitfalls that multi linage EMM types can cause for outbreak analysis in single reference based contexts.

### LIN code suitability for outbreak cluster detection

To assess the suitability of LIN codes for outbreak analysis, isolates from 5 clonal and 5 non-clonal EMM types were randomly selected from the UKHSA samples submitted to PubMLST, which had also previously undergone analysis as part of an outbreak investigation. This validation assessed whether LIN codes would infer the same outbreak groupings as SNP-based distance matrices. Assessment was performed by evaluating the correlation between sets of isolates per outbreak, for SNP if 10 or fewer SNPs apart, and LIN code if grouped within threshold 6 (cutoff of 5-10 allelic differences). The outcomes showed that 8 out of 10 EMM types had 0 mismatches between each isolates LIN and SNP based inclusion within specific outbreak clusters (Table 4), with the remaining 2 having 1 mismatched isolate by cluster assignment. For EMM1, the mismatches occurred where a LIN code difference at threshold 6 was present, indicating more than 5 but less than 10 allelic differences, when 9 SNP differences were detected. For EMM108, there were 14 SNP differences between clusters, but only differed at the 7^th^ LIN code threshold, which indicate a difference of 3-5 alleles. The high level of concordance shown across EMM types by SNP distance and LIN code show an appropriate level of cluster matching, with finer grained LIN code levels allowing for more granular definition and ascertainment of intra outbreak isolate relatedness.

**Table 4.** Table outlining correlation of cluster membership at LIN code threshold 6 (5-10 allelic differences), and 10 or fewer SNP differences by EMM type and number of representative isolates. In each instance mismatching was between 0 and 1 isolates indicating a high level of correlation.

| EMM type | Isolates (n) | Cluster mismatch 5-10 alleles LIN 6 (%) | Reason for differences |
| --- | --- | --- | --- |
| EMM1 | 16 | 6.3 | SNP difference of 9 from rest of cluster, difference at LIN threshold 6. |
| EMM3 | 32 | 0 |  |
| EMM8 | 49 | 0 |  |
| EMM11 | 138 | 0 |  |
| EMM28 | 27 | 0 |  |
| EMM33 | 48 | 0 |  |
| EMM59 | 35 | 0 |  |
| EMM77 | 50 | 0 |  |
| EMM82 | 7 | 0 |  |
| EMM108 | 22 | 4.5 | 14 SNPs from cluster, 2-5 allelic differences by LIN code |

A selection (n=9) of EMM77 outbreaks, previously analysed by UKHSA, were selected for further investigation as to the suitability of the *S. pyogenes* LIN coding scheme (Figure 7). A SNP based phylogenetic tree was produced by Gubbins (Figure 7), with branch lengths representing the logarithm of SNP distance to enable better visualisation of the EMM77 isolates, annotated by investigation and with LIN Codes. EMM77 was chosen due to its multi-lineage nature to ensure that the LIN coding scheme can effectively delineate distant lineages and outbreak relevant distances.

**Figure 7.**
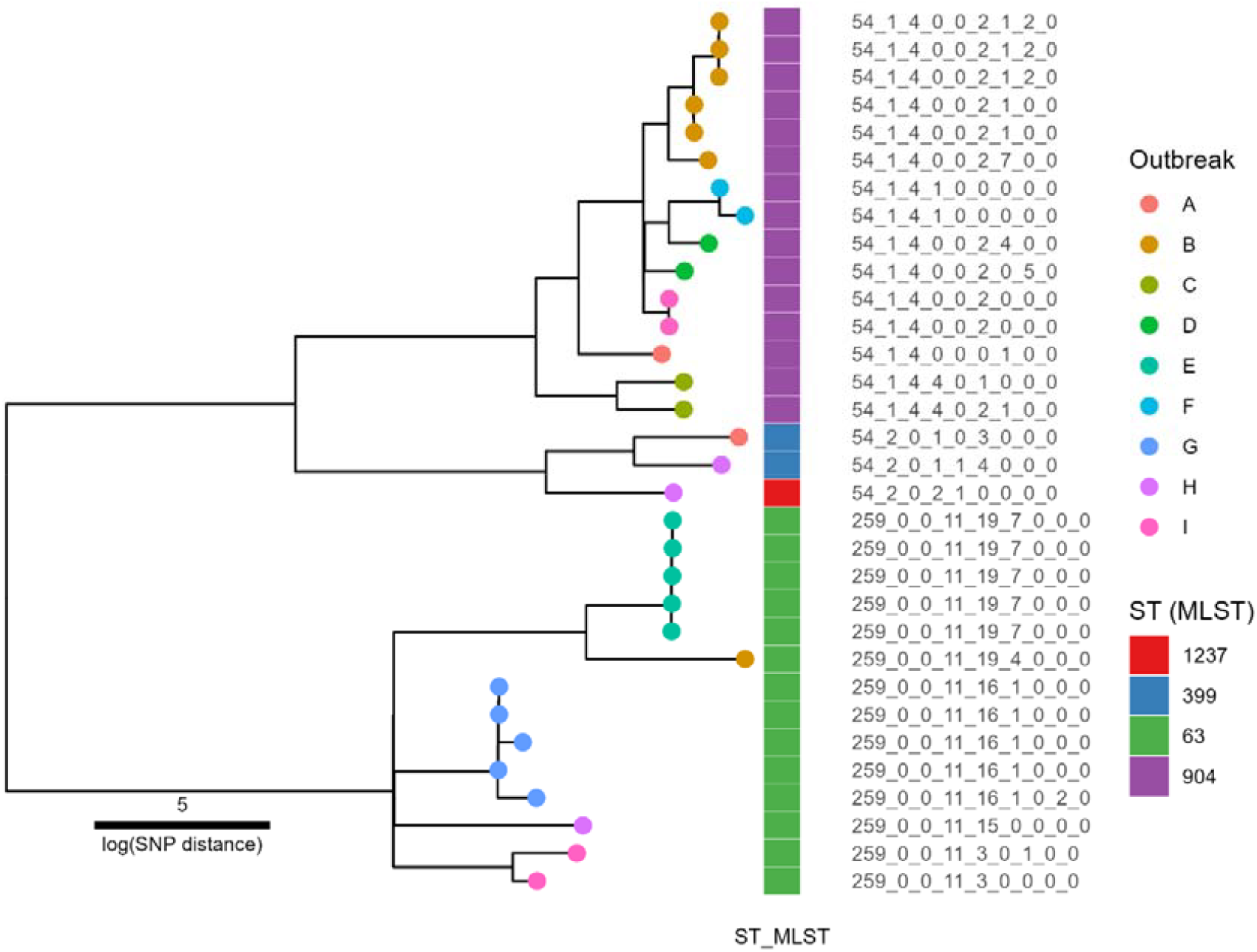
SNP based phylogenetic tree created using the PHEnix pipeline, and ggtree in R. The tree has been labelled with LIN codes as tips, with the colour of the circles representing individual outbreaks, and annotated with MLST ST. The phylogenetic tree showing two clades of EMM77, annotated with colours of their related outbreaks, annotated with their LIN codes. Each outbreak represents a separate investigation undertaken by UKHSA. LIN codes align well with the tree structure, and additionally highlight isolates which appear closely related via both SNP and LIN code across outbreaks.

The structure of the phylogenetic tree concurs with LIN code levels 1 and 2, with 54_1 (MLST ST904) and 54_2 (MLST ST’s 399 and 1237) matching the two subclades of the first clade (∼600 SNP differences), and with the second clade aligning fully with LIN code 259 (MLST ST 63) having ∼7500 SNPs difference. LIN codes for outbreak A correctly delineate by subclade, with the two isolates being ∼600 SNPs distant and by LIN code are between 528 and 285 allelic differences. Outbreak B is likewise correctly matched in 6 out of 7 isolates until the sixth threshold, with a range of 0-5 allelic differences, and over 528 allelic differences to the ST63 isolate. Similar patterns are shown across the other outbreaks, with isolates with 0 branch lengths producing indistinguishable LIN codes (e.g. outbreak E), and those with multi-lineage strains present (e.g. outbreak I) LIN codes which correctly reflect this.

## Discussion

There is a pressing global need for scalable, high-definition population biology study of *S. pyogenes*, due to its commonality, increasing prevalence, cost to human health and challenge in identifying clusters due to temporal and geographic spread (Nabarro, Brown et al. 2022, Smith, Marchant et al. 2025). This study found that our novel *S. pyogenes* cgMLST and LIN coding schemes can be used to facilitate epidemiological analysis for outbreak control and population level study. Hosting on PubMLST makes this a free and open resource, enabling international collaboration with a single mutually understandable nomenclature.

Healthcare systems across the world face challenges from GAS with low and high income nations along with differing geographies carrying different combinations of circulating strains (Smeesters, de Crombrugghe et al. 2024), resulting in different clinical requirements, resources for treatment and in investigating and controlling outbreaks.

The latest methods for assessing the genetic relatedness of *S. pyogenes* isolates use WGS derived SNP distances (Sharma, Ong et al. 2019) or cgMLST (Friães, Mamede et al. 2022, Toorop, Kraakman et al. 2023). SNP based and cgMLST based methodologies provide the highest levels of resolution, but SNP based approaches require rerunning of analyses as new isolates are investigated, and if used in conjunction with tools such as snapperDB are limited by reference strain and unstable nomenclature that is only comparable within that snapperDB instance (Dallman, Ashton et al. 2018). cgMLST schemes are typically hosted online, but in the case of the Toroop scheme, only those who have purchased the Ridom SeqSphere software can submit new alleles for inclusion in the scheme. PubMLST meanwhile allows for anyone to submit new alleles or genome for allele calling and assignment, thus enables seamless collaboration and mutually comparable results.

Since 2023, UKHSA sequenced on average 4700 *S. pyogenes* isolates a year, the analysis of which resulted in significant computational time and cost. Iteratively re-running pairwise SNP based analysis for more common EMM types becomes quickly infeasible without pruning isolates in the analysis, limiting the ability for national comparison and lookbacks over time. cgMLST methods meanwhile have the benefit of not requiring compute intensive pairwise analysis, require a single set of references rather than EMM type specific references. cgSTs or hashed cgMLST profiles can additionally be used to compare isolates between labs with greater ease (Deneke, Uelze et al. 2021). cgMLST allelic profiles were used to generate a minimum spanning tree of all uploaded isolates for this investigation, annotated with EMM type, MLST ST and level 0 LIN code for *S. pyogenes* showed strong correlation between assignments. The major and expected differences occurred in multi-lineage EMM types which occurred on different branches and matched to different ST’s and LIN0 assignments (which however did correlate). Parity between these different typing methodologies shows that these differences are effectively captured within the cgMLST scheme.

LIN coding provides a mechanism to utilise the benefits of cgMLST with a stable classification system, without risk of merges or alterations on the addition of new isolates. The LIN coding scheme as set out in this publication utilises 9 thresholds, covering the range from broader species wide genomic structure correlating with EMM and MLST ST to granular allelic differences for outbreak analysis (Pearce, Alikhan et al. 2018). LIN codes in this analysis (Figure 7) have been shown to be able to capture broader genomic structure with multi-lineage EMM types alongside the ability to aid outbreak investigations via exclusion of isolates too distant to be related, and identifying which isolates are potentially epidemiologically related and to what degree. Testing across 10 different EMM types (Table 4.) compounded these results, with an average concordance of 99% between a 10 SNP cutoff, and 10 allelic difference threshold definition of inclusion as an outbreak cluster. Compared to SNP based genetic relatedness methods which require a reference per EMM type and significant bioinformatician and computational time, isolates uploaded to PubMLST will automatically undergo cgMLST allele calling; if the cgST has been previously assigned a LIN code will immediately be assigned, if novel, the process to assign new LIN code will be run nightly and then be assigned. Due to being hosted on PubMLST, international and interlaboratory comparisons are easily enabled by sharing isolate ID’s or LIN codes, owing to their mutual comprehensibility without the need to transfer sequence data between parties (Jolley, Bray et al. 2018).

For a high throughput lab such as the Staphylococcus and Streptococcus reference section to undertake national surveillance, SNP based analysis is not economically able to systematically compare each newly sequenced isolate to all prospectively related isolates. LIN codes in comparison, enable long term national surveillance and real time monitoring of all newly sequenced isolates. For ongoing investigations, newly sequenced isolate’s LIN codes can be readily compared to other isolates within a cluster, providing an easy and effective method to ascertain the relatedness of isolates in an outbreak. This can be taken further to allow automated comparison of newly assigned LIN codes against previous assignments, to identify cryptic clusters and alert the relevant parties to an outbreak. In instances where a novel strain is discovered with potential international spread (Beres, Olsen et al. 2024, Vieira, Wan et al. 2024, Davies, de Gier et al. 2025), LIN codes can quickly be shared and used to determine potential cross border transmissions and inform public health measures.

The newly developed cgMLST and LIN coding scheme for *S. pyogenes*, are hosted on PubMLST and publicly available. The scheme represents a step forward in the global collaborative effort for surveillance and genetic relatedness assessment of *S. pyogenes*, providing a multi-resolution approach, which can be used for facilitating global public health.

## Acknowledgements

Thank you to the UKHSA Staphylococcus and Streptococcus Reference Section lab team for their tireless work which greatly benefited this project.

With thanks to Charlene Rodrigues who made the introductions to enable this collaboration and work between UKHSA and the University of Oxford.

